# Severe Biallelic Loss-of-function Mutations in *Nicotinamide Mononucleotide Adenylyltransferase 2 (NMNAT2)* in Two Fetuses with Fetal Akinesia Deformation Sequence

**DOI:** 10.1101/610899

**Authors:** Marshall Lukacs, Jonathan Gilley, Yi Zhu, Giuseppe Orsomando, Carlo Angeletti, Jiaqi Liu, Xiuna Yang, Joun Park, Robert J. Hopkin, Michael P. Coleman, R. Grace Zhai, Rolf W. Stottmann

**Affiliations:** Divisions of Human Genetics and Cincinnati Children’s Hospital Medical Center, Department of Pediatrics, University of Cincinnati, Cincinnati, OH, 45229, USA; John van Geest Centre for Brain Repair, University of Cambridge, ED Adrian Building, Forvie Site, Robinson Way, Cambridge, CB2 0PY, UK; Signalling ISPG, The Babraham Institute, Babraham, Cambridge CB22 3AT, UK; Department of Molecular and Cellular Pharmacology, University of Miami Miller School of Medicine, Miami, FL 33136, US; Department of Clinical Sciences (DISCO), Section of Biochemistry, Polytechnic University of Marche, Via Ranieri 67, 60131, Ancona, Italy; School of Pharmacy, Key Laboratory of Molecular Pharmacology and Drug Evaluation (Yantai University), Ministry of Education, Collaborative Innovation Center of Advanced Drug Delivery System and Biotech Drugs in Universities of Shandong, Yantai University, Yantai, Shandong 264005, China; Developmental Biology, Cincinnati Children’s Hospital Medical CenterDepartment of Pediatrics, University of Cincinnati, Cincinnati, OH, 45229, USA

## Abstract

The three nicotinamide mononucleotide adenylyltransferase (NMNAT) family members synthesize the electron carrier nicotinamide adenine dinucleotide (NAD^+^) and are essential for cellular metabolism. In mammalian axons, NMNAT activity appears to be required for axon survival and is predominantly provided by NMNAT2. NMNAT2 has recently been shown to also function as a chaperone to aid in the refolding of misfolded proteins. *Nmnat2* deficiency in mice, or in its ortholog *dNmnat* in *Drosophila*, results in axon outgrowth and survival defects. Peripheral nerve axons in NMNAT2-deficient mice fail to extend and innervate targets, and skeletal muscle is severely underdeveloped. In addition, removing NMNAT2 from established axons initiates axon death by Wallerian degeneration. We report here on two stillborn siblings with fetal akinesia deformation sequence (FADS), severely reduced skeletal muscle mass and hydrops fetalis. Clinical exome sequencing identified compound heterozygous *NMNAT2* variant alleles in both cases. Both protein variants are incapable of supporting axon survival in mouse primary neuron cultures when overexpressed. *In vitro* assays demonstrate altered protein stability and/or defects in NAD^+^ synthesis and chaperone functions. Thus, both patient *NMNAT2* alleles are null or severely hypo-morphic. These data indicate a previously unknown role for *NMNAT2* in human neurological development and provide the first direct molecular evidence to support the involvement of Wallerian degeneration in a human axonal disorder.

## INTRODUCTION

Fetal Akinesia Deformation Sequence (FADS) defines a broad range of disorders unified by absent fetal movement resulting in secondary defects often leading to stillbirth or limited postnatal survival ^1; 2^. These secondary features include edema, hydrops fetalis, craniofacial anomalies including micrognathia, lung hypoplasia, rocker bottom feet, intrauterine growth restriction, and decreased muscle mass ^3^. Through previous experimental models of fetal paralysis, the secondary findings have been shown to be primarily caused by a lack of fetal movement ^1; 4^. FADS has both genetic and environmental causes that can affect any aspect of the motor system including the central nervous system (CNS), peripheral nervous system (PNS), neuromuscular junction (NMJ), and/or skeletal muscle. Although most cases of FADS do not have a genetic diagnosis, multiple monogenic causes of FADS affecting PNS innervation development have been identified to date including *RAPSN, DOK7, MUSK* ^5–7^.

Through whole exome sequencing and subsequent Sanger sequencing of a family with two fetuses with FADS, we identified compound heterozygous mutations in a gene previously unlinked to FADS, *nicotinamide mononucleotide adenylyltransferase 2 (NMNAT2)*. NMNAT family members were first shown to play a role in axon degeneration with the discovery of the slow *Wallerian Degeneration (Wld*^*S*^) mutant mouse that showed delayed axon degeneration post transection ^8^. The *Wld*^*S*^ phenotype arose as the result of a spontaneous genomic rearrangement generating a fusion protein of NMNAT1 and the N-terminus of UBE4B, an E4 type ubiquitin ligase^9; 10^. Normally NMNAT1 is located only in the nucleus but the partial axonal location of the fusion protein leads to a gain-of-function explaining the slow Wallerian degeneration phenotype ^11^.

There are three canonical NMNAT isoforms and each displays unique subcellular localization and tissue specific expression. NMNAT1 is nuclear and broadly expressed, NMNAT2 is in the cytoplasm and axoplasm and enriched in the brain. NMNAT3 is proposed to be localized to the mitochondria and has lower expression in the brain ^12; 13^. The functions of all three have been studied in mice but until now only *NMNAT1* has been linked to human disease. *NMNAT1* mutations cause Leber’s Congenital Amaurosis 9 (LCA9) characterized by photoreceptor-neuron degeneration resulting in congenital blindness ^14–17^. Two N-ethyl-N-nitrosourea generated *Nmnat1* missense mouse mutants develop photoreceptor degeneration and closely model the pathology observed in LCA9 ^18^. In contrast, *Nmnat3* homozygous null mice show no nervous system phenotype and instead develop splenomegaly and hemolytic anemia ^19^. Hikosaka, et. al. showed NMNAT3 is the predominant NAD producer in the cytoplasm of mature erythrocytes and loss of *Nmnat3* resulted in defective glycolysis in these cells ^19^. To date, no patients with mutations in *NMNAT3* have been identified. These data illuminate the tissue specific requirements for NMNAT family members during development.

An essential role for NMNAT2 in axon growth and survival was established first by RNAi in primary neuronal culture and subsequently in *MNMAT2-deficient* mice ^20–22^. Acute removal of NMNAT2 *in vitro* from established axons causes axon degeneration through the Wallerian pathway, while its constitutive deletion in mice causes defects in PNS and CNS axon outgrowth, and consequent underdevelopment of the skeletal muscle which lacks innervation ^21; 22^. Other features shared with FADS include craniofacial defects and perinatal lethality due to a failure to inflate the lungs at birth ^21; 22^. Furthermore, RNAi of the *Drosophila* ortholog d*NMNAT* is also sufficient to trigger spontaneous degeneration of established axons and genetic mutation causes growth and survival defects in axons and their presynaptic termini ^23–25^. Conversely, overexpression of NMNATs after peripheral nerve transection can delay Wallerian degeneration and rescues all mouse NMNAT2 and *Drosophila* dNMNAT genetic knockdown or deletion phenotypes mentioned above ^13^. Deletion of another Wallerian pathway gene, *Sarm1*, also rescues axonal phenotypes in *MNMAT2-deficient* mice preventing perinatal lethality and allowing survival into old age with no overt behavioral changes ^26; 27^. These data demonstrate NMNAT2 protects against an active Wallerian Degeneration pathway mediated by SARM1. Recently, it has been discovered that NMNATs including NMNAT2 act as chaperones for protein refolding as well as NAD-synthesizing enzymes ^24; 28; 29^. *NMNAT2* transcripts have been shown to be decreased in human neurodegenerative diseases and the chaperone function of NMNAT2 has been shown to protect against neurodegeneration in a variety of tauopathy models ^29; 30^. While it remains controversial which function(s) of *NMNAT2* are neuroprotective, we sought to investigate both functions in our patient variants of *NMNAT2*. Interestingly, we found both functions are impaired. This finding and the striking similarity to the homozygous null mouse phenotype strongly support a causative role for these mutations.

## MATERIALS AND METHODS

### Subjects

Initial exome analysis was performed as a clinical service (Ambry Genetics). Informed consent to study the sequence data on a research basis was obtained according to Cincinnati Children’s Hospital Medical Center (CCHMC) institutional review board protocol # 2014-3789. Following consent, residual DNA samples were obtained for Sanger sequencing confirmation of exome sequencing analysis.

### Sanger Sequencing

Sanger sequencing to confirm the results of whole exome sequencing was performed by PCR amplification of exon 5 of *NMNAT2* with F primer 5’-gaggttcaggagcgatgaaa-3’ and R primer 5’-caggagaagagtgcacacca-3’ using genomic DNA. Exon 9 of *NMNAT2* was PCR amplified from genomic DNA with F primer 5’-gctcaaatgtgcttgctgaa-3’ and R primer 5’-cagacatgggatggttgatg-3’. Conservation and protein prediction scores for *NMNAT2* R232Q variant were generated by SIFT, Polyphen, and MutationTaster algorithms. Schematic of NMNAT2 protein domains were generated from existing literature and functional domains annotated by UniProt by homology. The crystal structure of NMNAT1 (PDB ID: 1kku) in stereo ribbon view was generated by PyMOL (v2.2.3)^31^.

### Histology

Histology was performed on formalin-fixed, paraffin-embedded patient tissue collected at the time of autopsy. Hematoxylin and eosin staining was performed according to standard methods at the CCHMC Pathology Core Lab and analyzed by attending pathology physicians.

### Constructs

The R232Q and Q135Pfs*44 *NMNAT2* mutations were introduced separately by QuikChangeII site-directed mutagenesis (Stratagene) into the complete open reading frame of the canonical 307 amino acid human NMNAT2 isoform cloned into expression vector pCMV-Tag2 (Stratagene). The expressed NMNAT2 proteins have a Flag tag and short linker sequence (17 amino acids) at their N-terminus. The presence of the mutations and absence of other PCR errors was confirmed by sequencing (Cogenics). pDsRed2-N1 (Clontech) was used for expression of variant *Discosoma* red fluorescent protein (DsRed) to label micro-injected neurons / neurites. pEGFP-C1 (Clontech) was used for expression of enhanced green fluorescent protein (GFP) to act as a transfection control and reference for NMNAT2 turnover in HEK 293T cells.

#### HEK 293T transfection and stability assays

HEK 293T cells were cultured in DMEM with 4,500 mg/L glucose and 110 mg/L sodium pyruvate (PAA), supplemented with 2 mM glutamine and 1% penicillin/streptomycin (both Invitrogen), and 10% fetal bovine serum. Cells were plated in 24-well format to reach 50-60% confluence for transfection with Lipofectamine 2000 reagent (Invitrogen) according to the manufacturer’s instructions. In standard turnover experiments (Fig. 4A) 500 ng Flag-NMNAT2 expression construct (wild-type or mutant), 200 ng of an empty pCMV-Tag series vector, and 100 ng pEGFP were transfected per well. To boost expression of the NMNAT2^Q135Pfs*44^ mutant 700 ng Flag-NMNAT2 expression construct and 100 ng pEGFP-C1 were transfected per well (Fig. 4C). After treatment ±10 µM emetine hydrochloride (Sigma-Aldrich), cells from single wells were lysed directly in 100 µl 2x Laemmli sample buffer and heated to 100°C for 5 mins. Equal amounts of extract (either 10 or 15 µl) were resolved on 12% SDS polyacrylamide gels, transferred to Immobilon-P membrane (Millipore) and probed with antibodies essentially as described previously ^20^. The following primary antibodies were used: mouse monoclonal anti-FLAG M2 (1:2,000 Sigma-Aldrich F3165), mouse monoclonal anti-GFP clones 7.1 and 13.1 (1:2,000, Sigma-Aldrich 11814460001) and rabbit polyclonal α-Tubulin (1:7,500, Thermo Fisher Scientific PA5-29444). Appropriate HRP-conjugated secondary antibodies were used for band detection with SuperSignal™ West Dura Extended Duration Substrate (Thermo Fisher Scientific) using an Alliance chemiluminescence imaging system (UVITEC Cambridge). Relative band intensities on captured digital images were determined (area under histogram peaks) using Fiji software (http://fiji.sc) ^32^.

**Figure 4.**
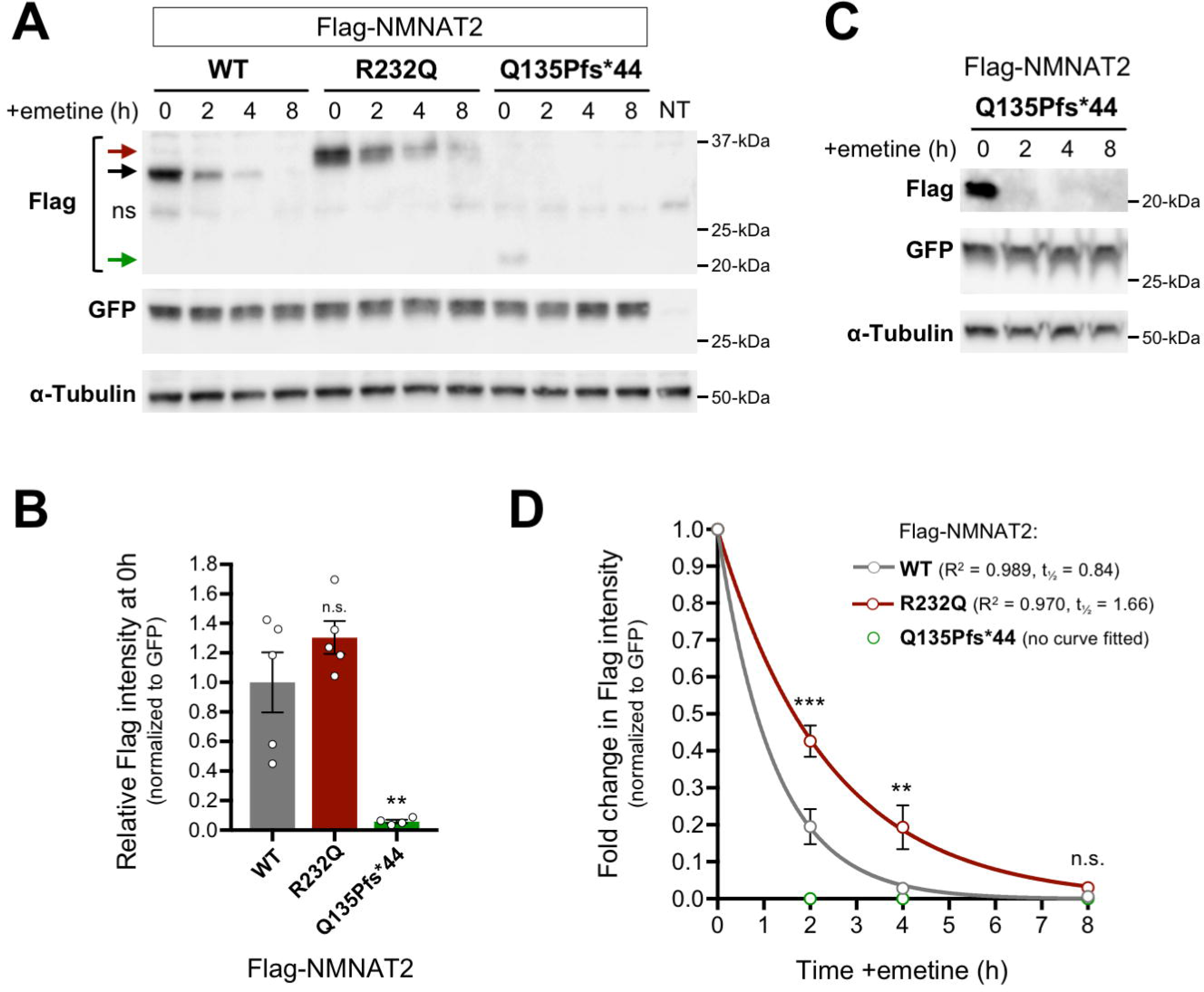
Relative stabilities and activities of NMNAT2^R232Q^ and NMNAT2^Q135Pfs*44^ in HEK 293T cells. (A) Representative immunoblots (of n = 3) of extracts of HEK 293T cells co-transfected with expression vectors for Flag-NMNAT2^WT^, Flag-NMNAT2^R232Q^ or Flag-NMNAT2^Q135Pfs*44^ and eGFP at the indicated times after addition of 10 µM emetine. Emetine was added 24 h after transfection. Extract from non-transfected cells is also shown (NT). Blots were probed with Flag, eGFP and α-Tubulin antibodies. To avoid saturation of the protein degradation machinery that might artificially slow rates of turnover, expression of the Flag-NMNAT2 proteins was kept relatively low by including empty vector as part of the transfection mix (see Materials and Methods). Co-transfected eGFP or endogenous α-Tubulin (present in transfected and non-transfected cells) are both relatively stable proteins and were respectively used as a reference for Flag-NMNAT2 protein turnover (to control for transfection efficiency) and for loading. Arrows indicate the positions of bands corresponding to Flag-NMNAT2^WT^ (black, ∼34 kDa), Flag-NMNAT2^R232Q^ (red, ∼37 kDa), and Flag-NMNAT2^Q135Pfs*44^ (green, ∼22 kDa). An asterisk indicates the position of a non-specific band. (B) Relative steady-state Flag-NMNAT2 protein band intensities (0h, just before emetine addition) after normalization to co-transfected eGFP for blots described in panel A. Individual values (n = 4-5) and means ±SEM are plotted. n.s. = not significant (p > 0.05), ** p < 0.01, one-way ANOVA with Tukey’s multiple comparisons test (only comparisons to Flag-NMNAT2^WT^. (C) Representative immunoblots (of n = 4) of extracts of HEK 293T cells, as in panel A, but transfected with a higher concentration of Flag-NMNAT2^Q135Pfs*44^ expression vector and with increased loading per lane to maximize Flag band intensity at the 0h time point so that its level at 0h is similar to that of Flag-NMNAT2^WT^ in panel A. This allows for a more accurate comparison of turnover rates. (D) Relative turnover rates of Flag-NMNAT2 proteins after emetine addition. Flag-NMNAT2 band intensities on blots described in panel A (Flag-NMNAT2^WT^ and Flag-NMNAT2^R232Q^) and panel C (Flag-NMNAT2^Q135Pfs*44^) were normalized to co-transfected eGFP and intensities at each time point after emetine addition were calculated as a proportion of the intensity of the 0h, untreated band. Means ± SEM (n = 4) are plotted. n.s. = not significant (p > 0.05), ** p < 0.01 and *** p < 0.001, two-way ANOVA with Sidak’s multiple comparisons test for effects between variants. One-phase decay curves were fitted to the data sets for Flag-NMNAT2^WT^ and Flag-NMNAT2^R232Q^ using non-linear regression. The R^2^ value and half-life (t½) are reported. No intensity values could be obtained for Flag-NMNAT2^Q135Pfs*44^ at any timepoint assessed after emetine addition precluding curve fitting and statistical analysis.

#### Microinjections and imaging

The preparation of dissociated SCG neuron cultures from wild-type P0-P2 pups, microinjections, Flag immunostaining and quantification of neurite survival were all performed essentially as described previously ^20; 22^. Expression vectors and the concentrations used in each specific injection experiment are described in the Figure 3 legend. Fluorescence images were acquired with a Leica DFC365FX fluorescence monochrome camera attached to a Leica DMi8 inverted fluorescence microscope (10x objective). Mean intensities of Flag immunostaining and DsRed fluorescence signals in injected SCG neurons were determined using Fiji software (http://fiji.sc) by thresholding (20, dark background) followed by particle analysis (size >250 pixels for 1392×1040 images) to identify neurons with signal intensity above background (the threshold value was subtracted from the mean intensity values obtained) ^32^.

**Figure 3.**
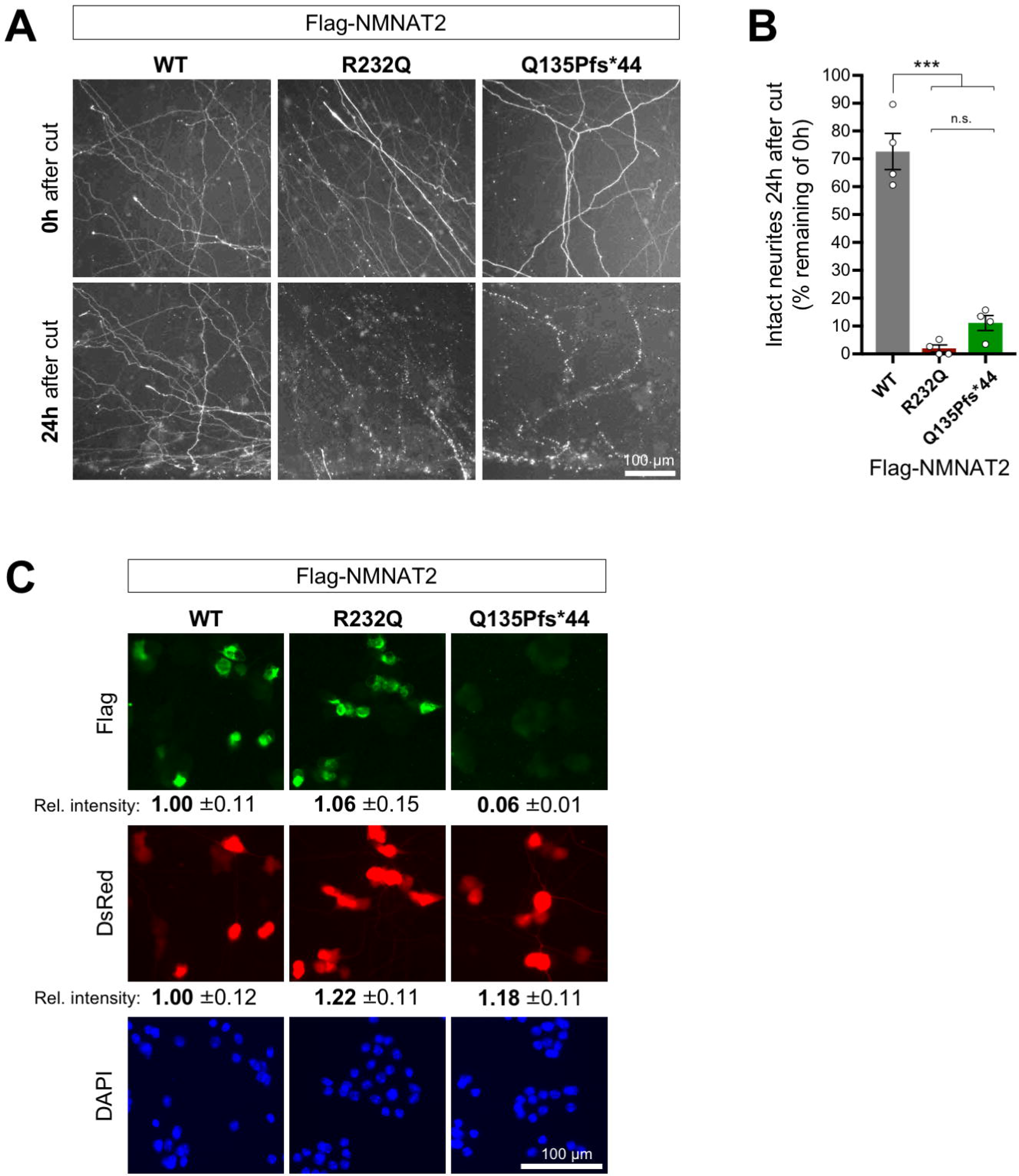
NMNAT2^R232Q^ and NMNAT2^Q135Pfs*44^ both have reduced capacity to maintain neurite survival. (A) Representative images (of n = 4 independent experiments) of cut neurites of SCG neurons co-injected with expression vectors for Flag-NMNAT2^WT^, or Flag-NMNAT2^R232Q,^ or NMNAT2^Q135Pfs*44^ (10 ng/µl) and DsRed (pDsRed2, 40 ng/µl). Neurites were cut 48 hours after injection when DsRed expression allows clear visualization of the distal neurites of the injected neurons. Images show transected neurites, just distal to the lesion, immediately after (0h) and 24 hours after cut. The lesion site is located the bottom edge of each field. Brightness and contrast have been adjusted for optimal visualization of neurites. (B) Quantification of neurite survival at 24 hours after cut for experiments described in panel A. The number of intact neurites with continuous DsRed fluorescence at 24 hours is shown as a percentage of intact neurites at 0h. Individual values and means ± SEM are plotted (individual values represent the average of two fields per separate culture). n.s. = not significant (p > 0.05), *** p < 0.001, one-way ANOVA with Tukey’s multiple comparisons test. (C) Relative expression level of Flag-NMNAT2 variants in injected SCG neuron cell bodies. Representative fluorescent images of SCG neurons 24 hours after co-injection with expression vectors for Flag-NMNAT2^WT^, Flag-NMNAT2^R232Q^ or Flag-NMNAT2^Q135Pfs*44^ and DsRed (each at 25 ng/µl). DsRed identifies injected neurons, Flag immunostaining shows expression of the Flag-NMNAT2 proteins, and DAPI labels nuclei. Relative intensities (± SEM) of Flag immunostaining and DsRed signal are shown after transformation to the mean of levels in neurons injected with the Flag-NMNAT2^WT^ construct. The data for WT, R232Q and Q135Pfs*44 were calculated from 47, 62 and 40 injected neurons (DsRed positive) of which 87.2%, 81,3% and 22.5% were Flag-positive respectively.

### NMNAT recombinant protein expression and purification

For biochemistry assays (Fig. 5A-G), pET28c plasmid constructs were generated for NMNAT2^R232Q^ and NMNAT2^Q135Pfs*44^ pET28c to produce recombinant proteins with an N-terminal His tag and linker (MGSSHHHHHHSSGLVPRGSH) for affinity purification that matched a previously generated NMNAT2^WT^ construct ^33^. Expression was carried out in *E. coli* BL21(D3) cells (Invitrogen) following 0.5 mM IPTG induction for 4 h at 25°C with subsequent purification using TALON chromatography (Clontech) as described ^34^. The purified proteins were desalted on PD-10 columns (GE Healthcare) in 50 mM HEPES/NaOH buffer, pH 7.5, 1 mM Tris(2-carboxyethyl)phosphine (TCEP), 20 % glycerol, and stored at −80 °C. Their amount was measured by the Bio-Rad protein assay. Their purity was evaluated on SDS polyacrylamide gels either after Coomassie staining or immunoblotting. Proteins were transferred from gels to Immobilon-P membrane (Millipore) and probed with antibodies as described ^20^. Monoclonal anti-NMNAT2 (1:1,000 Abcam AB5698) or anti-tetra His (0.1 μg/ml Qiagen 34670) were used as primary antibodies, followed by appropriate HRP-conjugated secondary antibodies. SuperSignal™ West Dura Extended Duration Substrate (Thermo Fisher Scientific) was used for detection on an Alliance chemiluminescence imaging system (UVITEC Cambridge).

**Figure 5.**
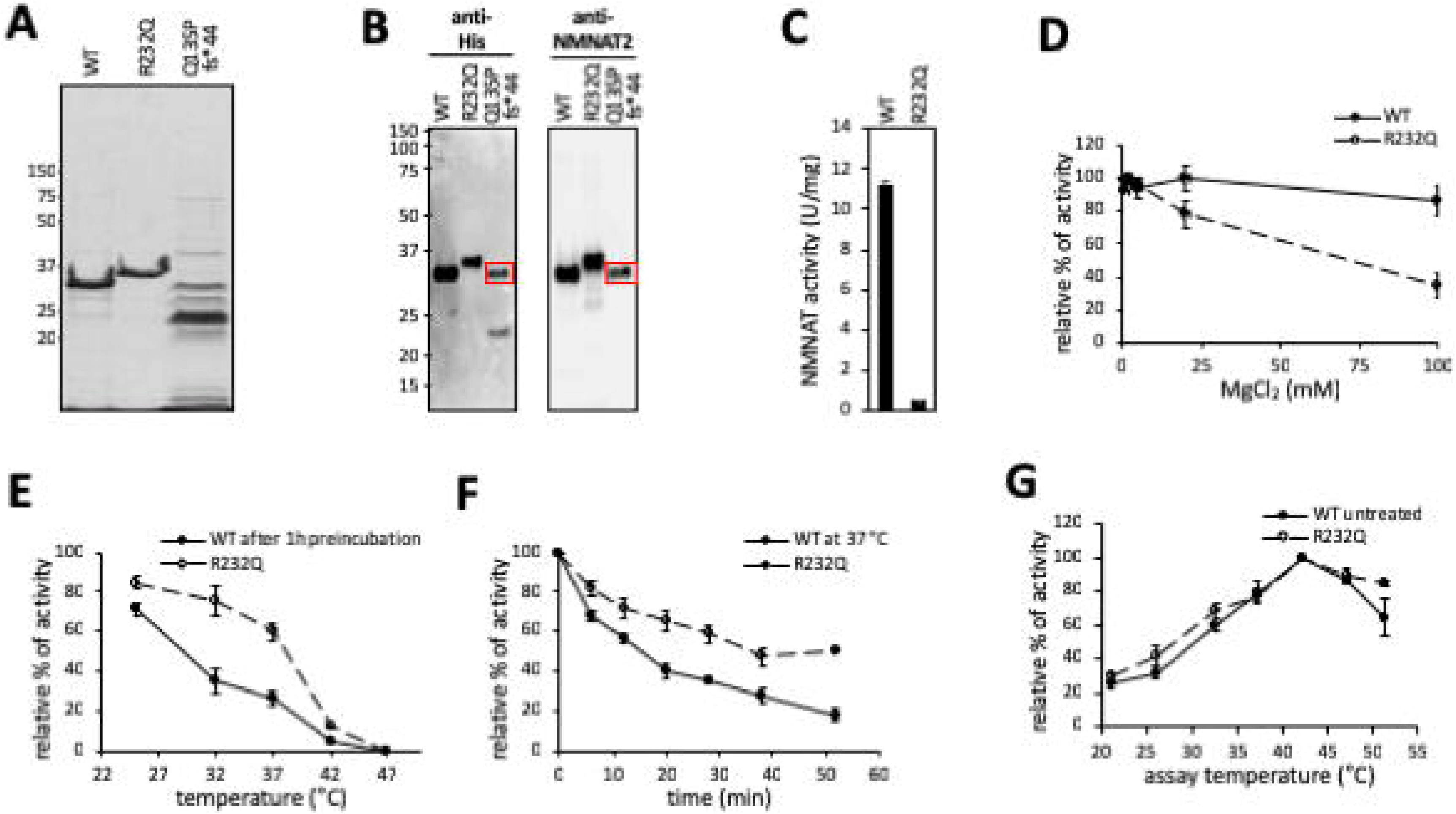
Bacterial expression and in vitro characterization of the activity of recombinant NMNAT2^R232Q^ and NMNAT2^Q135Pfs*44^. (A) Coomassie blue stained 12 % SDS polyacrylamide gel loaded with similar amounts (∼3 μg) of NMNAT2^WT^ and each indicated recombinant NMNAT2 variant arising from His-tag affinity chromatography. (B) Immunoblots of ∼0.3 µg of the same protein samples as in A probed with anti-His and anti-NMNAT2 antibodies as indicated. As in HEK cells, bacterially-expressed NMNAT2^R232Q^ migrates slower than NMNAT2^WT^ and NMNAT2^Q135Pfs*44^, which lacks the epitope recognized by the NMNAT2 antibody (raised against the C-terminus of the full-length protein), is expressed at a low level. The NMNAT2^Q135Pfs*44^ preparation contains a ∼34 kDa protein recognized by both anti-His and anti-NMNAT2 antibodies that is likely to be His-tagged NMNAT2^WT^ (red boxes). (C) NMNAT specific activity of His-tag purified preparations measured at 37 °C with saturating concentrations of substrates. The less pure NMNAT2^Q135Pfs*44^ preparation is omitted despite some activity found since it was not associated with the His-tagged 22 kDa truncated protein arising from the frame shift mutation (see text). (D) Magnesium-dependent rates of NMNAT activity referred to 1 mM MgCl_2_ (arbitrary 100 % value). (E) Enzyme stability after 1 hour treatment at different temperatures. Treated enzyme solutions were then assayed at 37 °C. Relative rates are expressed as percentages of the untreated enzyme kept at 4 °C (100 % not shown). (f) Enzyme stability at 37 °C as function of time. Rates are relative to time zero. (g) Optimum temperature after heating of whole assay mixtures at the indicated temperatures. Relative rates are expressed as percentages of the maximum observed (42 °C for both enzymes). All data presented are the mean ± SEM from n = 3 independent measures. T test p values *vs* corresponding WT are marked by (*) p < 0.015 or by (**) p < 0.005 (Two Sample t Test, unequal variances).

### NMNAT enzymatic activity assay

Routine assays were done by a spectrophotometric coupled method as described, in 0.5 mL mixtures containing 30 mM HEPES/NaOH buffer, pH 7.5, 0.5 mg/mL bovine serum albumin (BSA Sigma-Aldrich A7906), 75 mM ethanol, 30 mM semicarbazide (Sigma-Aldrich S2201), 12.5 U/mL alcohol dehydrogenase (ADH Sigma-Aldrich A7011) ^35^. NMNAT2^WT^ was assayed at 25 mM MgCl_2_ and 1 mM of both ATP and NMN (Fig. 5C). The mutant R232Q was assayed at 5 mM MgCl_2_, 5 mM Mg-ATP, and 1 mM NMN (Fig. 5D). For Mg^2+^-dependence studies the MgCl_2_ was increased up to 100 mM. Temperature studies were carried out under various treatments as indicated (Fig. 5E-G) using apo-enzyme solutions in buffer. With assays at 52°C, the reaction mixture at the end of incubation was cooled down to 37 °C and then enzyme was re-added to check for activity recovery, thus ruling out heat inactivation of the ancillary enzyme ADH. The *K*_m_ and *K*_cat_ values were calculated at 37 °C as described using 0.5-5 mM Mg-ATP and 0.05-1 mM NMN for the mutant R232Q, or 0.05-0.6 mM ATP and 0.01-0.15 mM NMN for the wild type ^36^. Due to the known instability of NMNAT2 preparations after thawing, enzyme was always added as the last component to start the reaction, and control assays were performed in parallel ^34^. One Unit (U) of NMNAT activity refers to the amount of enzyme that forms 1 μmol/min of product at the indicated temperature.

#### Gel filtration

Gel filtration of pure NMNAT2^R232Q^ was carried out by FPLC with a Superose 12 HR 10/30 column (Amersham Pharmacia), equilibrated with 50 mM HEPES/NaOH buffer, pH 7.5, 0.15 M NaCl, 1 mM DTT. Bovine serum albumin, ovalbumin, and carbonic anhydrase were used as the standards.

### In-cell luciferase refolding assay

HEK 293T cells were cultured in six-well plates and double transfected using jetPRIME transfection reagent (VWR International, Radnor, PA, USA) with pCMV-luciferase, and one of the following plasmids: pDsRed2 vector (control), pCMV-Hsp70, pCMV-Nmnat3, pCMV-Nmnat2^WT^, pCMV-NMNAT2^R232Q^, and pCMV-NMNAT2^Q135Pfs*44^. At 48 hrs after transfection, protein synthesis was inhibited by adding 1 μg/ml cycloheximide. Cells were subjected to heat shock at 42 °C for 45 mins, and then recovered at 37 °C for 3 hours ^37^. Cells were lysed in lysis buffer containing 100 mM KCl, 20 mM HEPES, 5% glycerol, 0.1% Triton X-100, and 1mM dithiothreitol. Luciferase activity was measured with the Luciferase Assay System (Promega, Madison, WI, USA).

#### Statistics

Statistical testing of data was performed using Excel (Microsoft) or Prism (GraphPad Software Inc., La Jolla, USA). The specific tests used are described in Figure legends.

## RESULTS

### Clinical Summary of two fetuses with FADS

#### Fetus 1

Fetus 1 was born to a 32yo Caucasian female evaluated for non-immune hydrops fetalis identified at 21 weeks gestation by ultrasonography. Ultrasonography identified multiple abnormalities including cystic hygroma, skin edema, ascites, and pleural effusion. Fetal MRI confirmed these findings and revealed profound hydrocephalus and cystic hygroma (Fig. 1A,B). The fetus was motionless though there was a normal amount of amniotic fluid. Amniocentesis showed a normal 46, XX karyotype and microarray analysis was negative for aneuploidy. Whole genome chromosome SNP microarray analysis was normal as well. Alpha fetoprotein was elevated and viral PCRs for toxoplasmosis, parvovirus, cytomegalovirus, and HSV were all negative.

**Figure 1.**
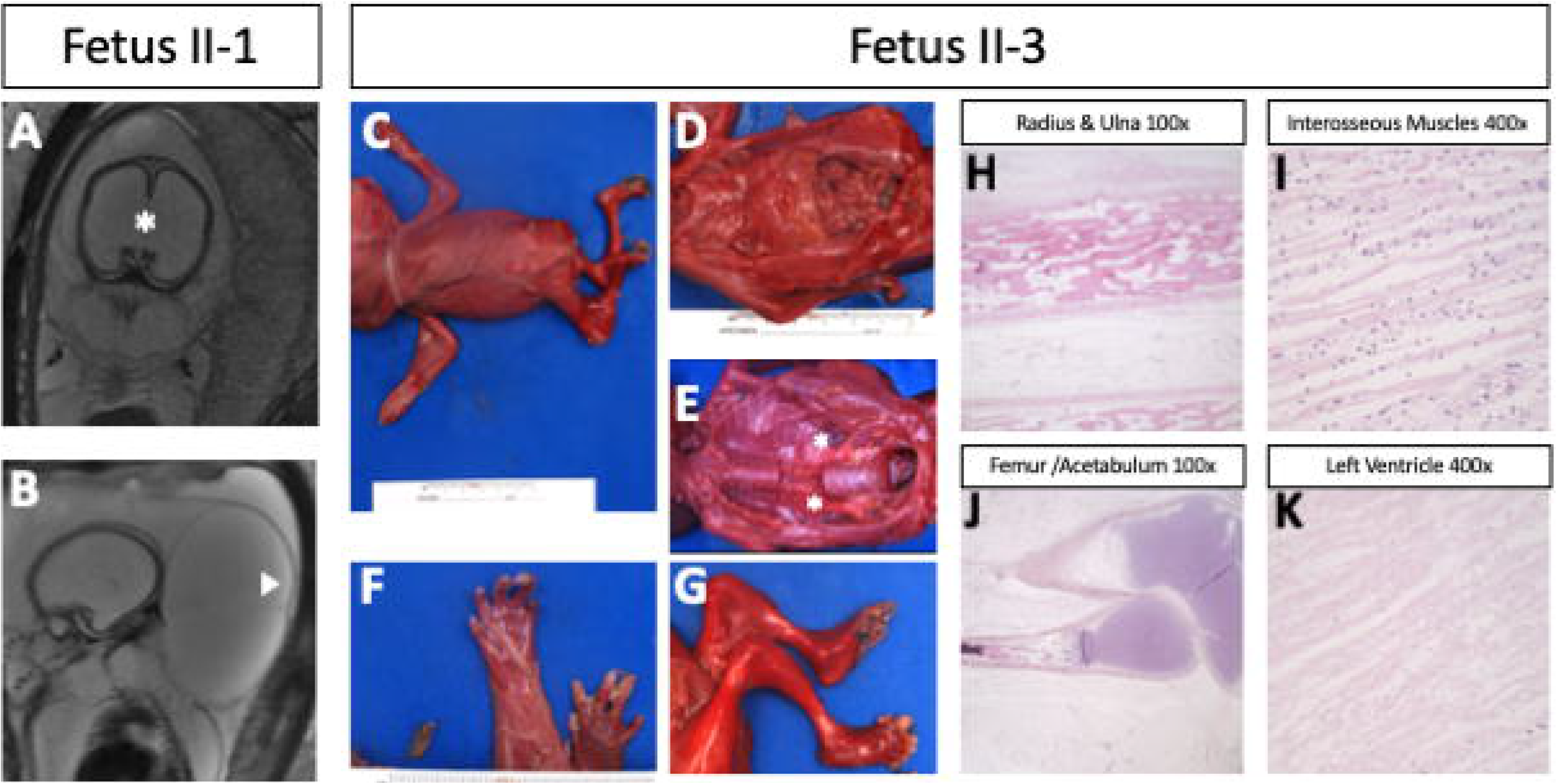
Gross phenotype and histology of affected fetuses. (A,B) Fetal MRI of Fetus II-1 in which hydrocephalus is noted by asterisk and cystic hygroma by arrowhead. (C) Dorsal view of fetus II-3 with notable edema and lack of skeletal muscle in the extremities. (D) Fetus II-3 displays malrotation of the gut with the appendix in the upper left quadrant. (E) Absence of the psoas muscles bilaterally is noted by the asterisks. (F,G) Fetus II-3 displays flattened hands with contractures of the elbow (F), and nearly complete absence of skeletal muscle of the leg (G). (H) Histology of the right radius and ulna shows reduced skeletal muscle fiber packing near the bone. (I) Histology of the interosseous muscles of the right hand show sparsely spaced muscle fibers with plump nuclei. (J) Histology of the hip joint shows fibrofatty tissue replacement of the musculature of the hip (K) Histology of the left ventricle shows normal architecture of the myocardium.

Fetal echocardiogram showed normal fetal cardiac anatomy, function, and rhythm approximately 2 weeks prior to delivery. Fetal MRI showed a head circumference that appeared to be within the normal range but showed evidence of severe dilation of the lateral and third ventricles (Fig.1A). The pons, cerebellum, and spinal cord were thin. Both kidneys were below the 5^th^ percentile in size for gestational age. Contrary to the findings in *MNMAT2-deficient* mice, there were no abnormalities in the bladder either grossly or microscopically ^21^. Both extremities remained in position and muscle planes were markedly diminished during fetal MRI. The fetus was delivered stillborn at approximately 27 weeks gestation.

Fetal autopsy was performed at Cincinnati Children’s Hospital Medical Center. Gross inspection identified multiple congenital anomalies including: hydrops fetalis, cystic hygroma, bilateral hypoplastic lungs, hydrocephalus, hypoplastic cerebellum, severely reduced skeletal muscle mass or absence, flexion contractures of all extremities, micrognathia, cleft palate, and hydropic placenta (Table 1). All tissues were extremely edematous with focal hemorrhage of soft tissues. The musculature was very poorly developed and nearly absent in all extremities. The lungs were hypoplastic. The placenta was grossly hydropic with evidence of chorioamnionitis and very friable, spongy, dark red tissue with no focal areas of discoloration or evidence of infarcts. The umbilical cord had an eccentric insertion and the fetal membranes were not discolored. The spinal cord was thin and poorly developed throughout most of its length. A frozen section of the quadriceps muscle showed no recognizable skeletal muscle tissue and appeared to be composed essentially of immature fat tissue. There were no inflammatory cell infiltrates or evidence of degenerating or dysplastic skeletal muscle fibers. The brain was poorly developed and collapsed when the calvaria was opened along the suture lines. Evaluation of the brain was severely limited due to autolytic changes, however, sections showed very immature neural glial tissue and germinal matrix. Only one slide showed a few areas of recognizable cortex.

**Table 1.**
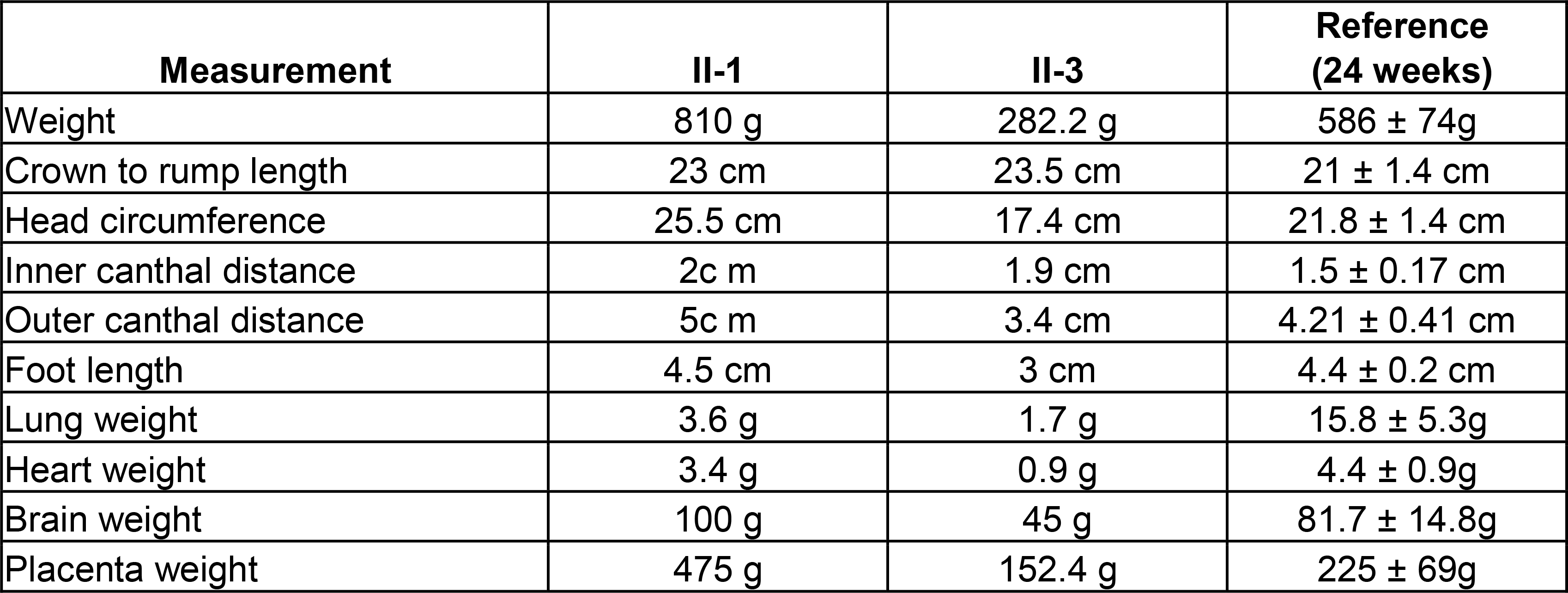
Clinical Features of fetuses II-1 and II-859 3

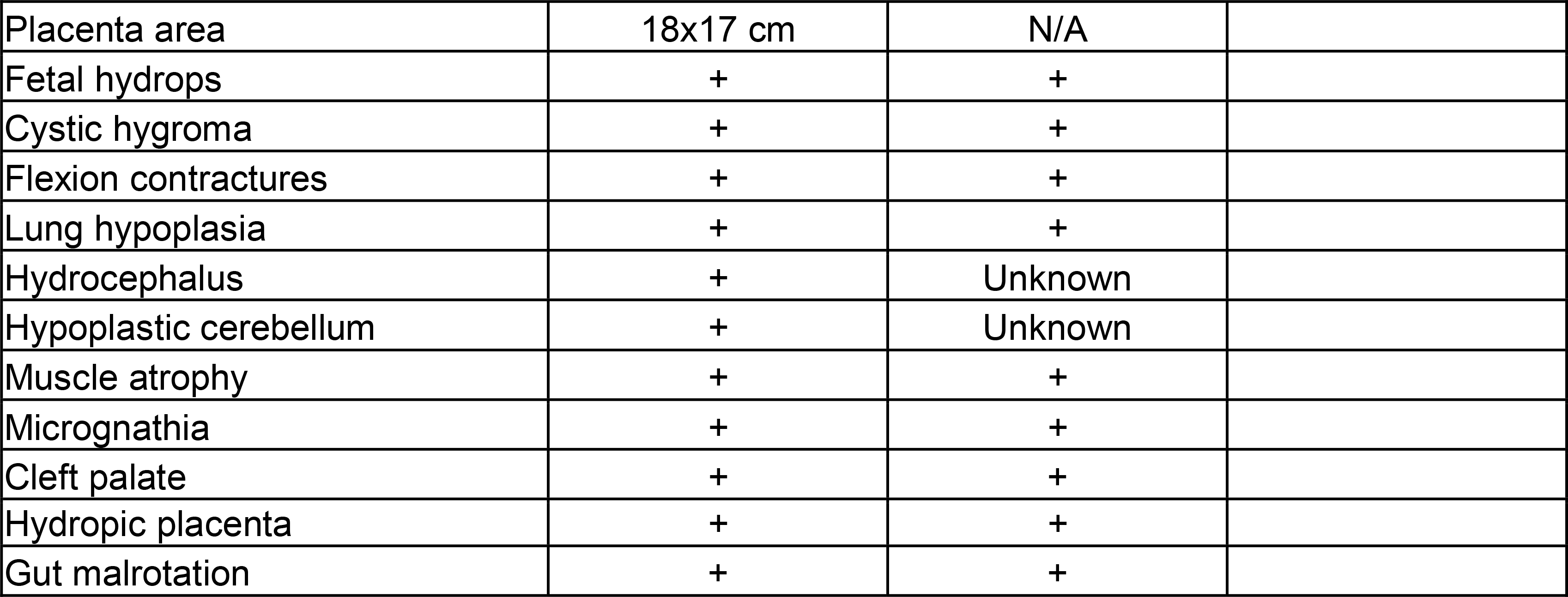

Given this constellation of phenotypes, the cause of death in fetus 1 was determined to be hydrops fetalis with multiple congenital anomalies and fetal akinesia deformation sequence (FADS). It is unlikely the phenotypes were due to a congenital myopathy based on evaluation of the muscle biopsy taken at autopsy.

#### Fetus 2

Fetus 2 was a second stillborn fetus of the same parents with an intervening healthy child (Fig.2A). The genetic analysis described below was used for prenatal counseling and fetus 2 was the product of prenatal genetic diagnosis/*in vitro* fertilization (PGD/IVF) and was intended to be a carrier for *NMNAT2* but developed severe hydrops at 16 weeks gestation as determined by Ultrasound (US). Given the prognosis of fetus 1, fetus 2 was delivered at 23 weeks gestation by Cesarean section and submitted for autopsy. Fetus 2 was diagnosed with hydrops fetalis with multiple fetal anomalies similar to fetus 1. Upon gross inspection, the fetus displayed hydropic changes including diffuse body wall and soft tissue edema with prominent nuchal fold (Fig. 1C). Markedly, there was apparent absence of skeletal muscle, especially that of the shoulder, extremities, pelvic girdle and absence of the psoas muscles, bilaterally (Fig. 1C-E). Long bone formation appeared adequate in length but slender with reduced formation of the femoral neck and trochanters. Sections of the limbs showed adequately formed long bones with adequate bony trabecular marrow space with scant trilineage hematopoiesis. The surrounding tissue was composed of mainly fibroadipose tissue in an edematous background with very few skeletal muscle fibers (Fig. 1H, J). The nuclei of the skeletal muscle were plump and evenly distributed at the edges of the pink proteinaceous myocyte fibers (Fig. 1I).The carpal cartilages were fused. Both the upper and lower limbs showed an abnormal absence of bundling of skeletal muscle fibers. The hands and feet had an unusual flattened appearance with severe contractures but adequate ray and digit bony formation (Fig. 1F,G). The heart was of normal weight with apparently normal myocardium, indicating the skeletal muscle was affected but the cardiac muscle was spared (Fig. 1K). The lungs were hypoplastic with normal lobar formation and no focal lesions or other abnormalities. No lesions were identified in the kidneys or bladder. The brain showed no significant gyration as expected for gestational age.

The placenta showed variable villous immaturity with irregular contours and occasional trophoblastic inclusions. There was a vascular distribution of fetal thrombotic vasculopathy. The cause of death for fetus 2 was likely related to placental insufficiency with placental immaturity, hydrops fetalis, and high-grade fetal thrombotic vasculopathy. Fetus 2 thus displayed many features of FADS including severely diminished muscle mass, pulmonary hypoplasia, joint contractures, and micrognathia.

### Exome Sequencing and Filtering

Genetic analysis was performed upon birth of fetus 1. Karyotyping and whole genome SNP microarray were normal suggesting a monogenic disorder. Clinical whole exome sequencing of the trio (Father, Mother, and Fetus 1) (Fig. 2A) was performed at Ambry Genetics. Mean coverage was 77.0, 105.6, and 86.7 reads per base for the father, mother and fetus, respectively. Filtering was performed by Ambry as detailed in Table S1. Briefly, multiple inheritance models were tested resulting in 26 potential gene candidates (with 40 total coding alterations) over all models. Manual review was performed for sequencing artifacts, known polymorphisms, additional artifacts and benign alterations.

**Figure 2.**
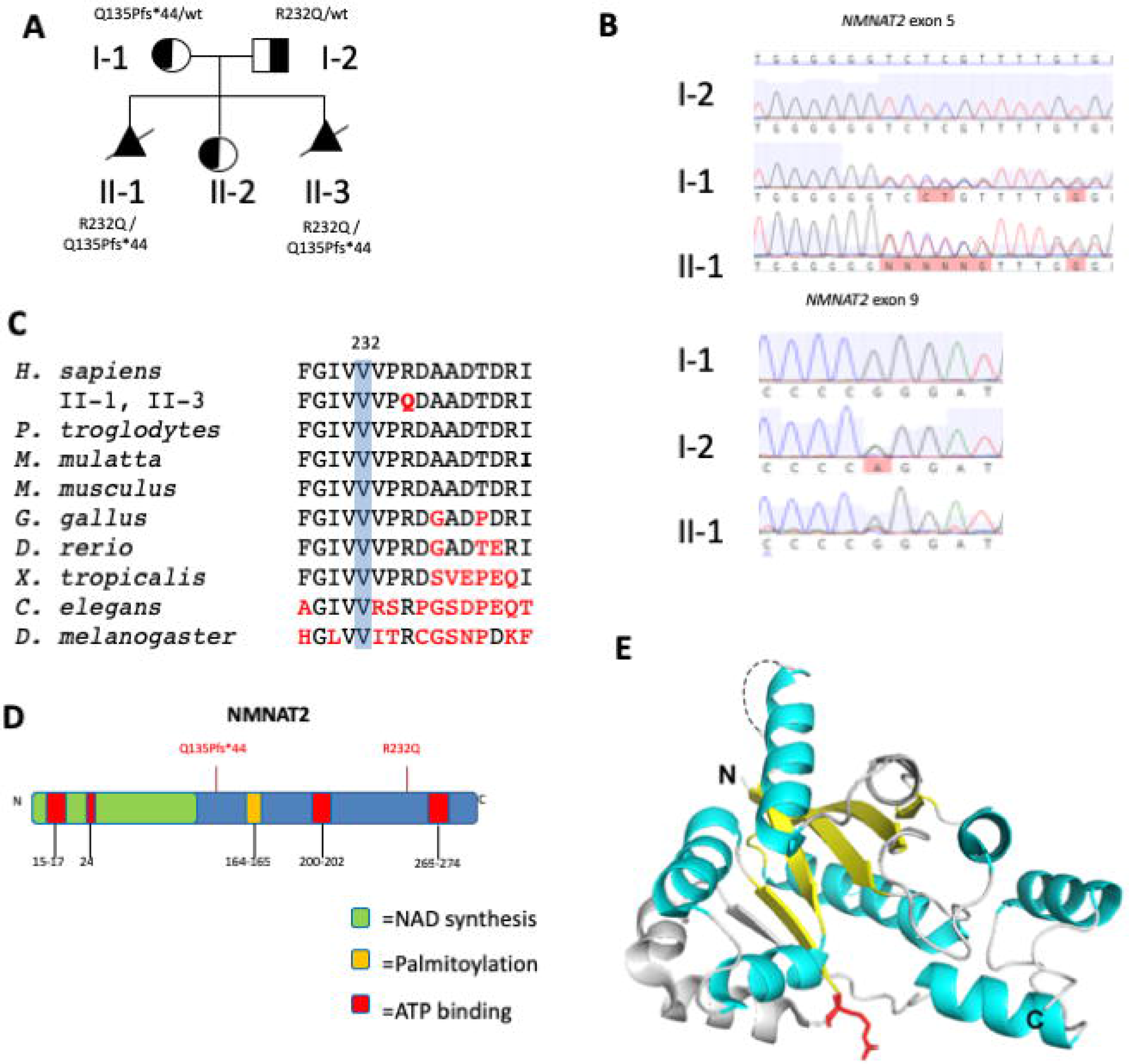
Whole exome sequencing identifies compound heterozygous mutations in *NMNAT2*. (A) Family pedigree. Stillborn infants are depicted as filled triangles with slashes (A). (B) Sanger sequencing of *NMNAT2*. (C) Conservation of arginine at aa232 in NMNAT2 homologues across distant phyla. (D) Diagram of functional domains of NMNAT2 with patient variant positions in red. (E) 3D structure model of NMNAT1. The conserved β strands and α helices important for enzymatic function are marked in yellow and cyan, respectively. The disordered region in NMNAT that contains the nuclear localization sequence is indicated by a dashed line. The R232 equivalent residue is labeled in red.

*NMNAT2* was identified as the remaining candidate in an autosomal recessive model with two unique alterations suggesting a compound heterozygous inheritance pattern (Supp Table 1). By co-segregation analysis, both parents were found to be heterozygous for one of the two mutations identified in fetus 1. *NMNAT2* was deemed clinically novel as no patients have been identified to date but there is significant phenotypic overlap between the patient’s phenotype with a mouse model of *NMNAT2* deficiency and the mutations are predicted to be damaging by SIFT and Polyphen ^21; 22^. When fetus 2 was diagnosed with such similar features, NMNAT2 was directly tested with Sanger sequencing and found to have the same compound heterozygous variants.

### *NMNAT2* Variants

The maternally inherited variant was a single duplication of a cytosine at position 403 in exon 5 resulting in a frameshift and premature stop after 44 amino acids in *NMNAT2* (c.403dupC, p.Q135Pfs*44; confirmed by Sanger Sequencing; Fig. 2B,D). The paternally inherited variant was a missense mutation in exon 9 (c.695G>A, p.R232Q; confirmed by Sanger Sequencing; Fig. 2B) resulting in a coding change of arginine to glutamine at position 232 (R232Q). Conservation alignment shows R232 is highly conserved to *D. melanogaster* and has a PhastCons score of 1 (Fig. 2C). Multiple algorithms predict this to be damaging sequence change (e.g.., PolyPhen score of 0.97, SIFT 0.0 and is predicted to be “disease causing” by MutationTaster. The family has one unaffected daughter who was identified to carry only one of the *NMNAT2* mutations. This finding supports a recessive model in which both affected alleles must be inherited in order to develop FADS.

Although the human NMNAT2 crystal structure has not been characterized, it is predicted that the enzyme activity domains share the same structure folds as those of human NMNAT1 and NMNAT3 ^38^. The R232 equivalent residue is invariant in all three human NMNAT isoforms and NMNAT homologs across distant phyla (Fig. 2C), suggesting its importance in protein function. Given that the R232 residue is located within the conserved region, we examined the crystal structure of human NMNAT1 to evaluate the potential structure-function consequences of the R232Q mutation in NMNAT2 ^39^ (Fig. 2E). The residue is located at the end of a β strand connecting the substrate binding domains for NMN and ATP, suggesting it is part of a conformational change upon binding the adenine group of ATP substrate or NAD/NaAD ^39^. It is predicted that the significant change in the side chain from arginine to glutamine will alter the electrostatic distribution of the substrate binding sites. In addition, R232 is at the bend between the β strand and a helix, positioned at the surface of the protein that likely participates in the interface of protein-protein interaction (Fig. 2E).

### Both the NMNAT2^R323Q^ and NMNAT2^Q135Pfs*44^ variants have reduced capacity to delay Wallerian Degeneration

Overexpression of mouse or human (Flag-tagged) NMNAT2 is sufficient to delay Wallerian degeneration in cultured mouse superior cervical ganglion (SCG) neurons. This affords us a ready assay to measure the capacity of a variant to support axon survival ^20; 40^. We therefore assessed whether this property is affected by the NMNAT2^R232Q^ or NMNAT2^Q135Pfs*44^ variants found in the affected patents. We introduced expression vectors for Flag-tagged wild-type or variant NMNAT2 into SCG neurons by microinjection. We used a concentration for which the resulting expression from the Flag-NMNAT2^WT^ construct preserves integrity of the majority (∼70%) of neurites of the injected neurons for at least 24 hours after transection. Under these conditions we found little or no preservation of cut neurites from SCG neurons injected with either Flag-NMNAT2^R232Q^ or Flag-NMNAT2^Q135Pfs*44^ expression vectors (Fig. 3A, B). This lack of protection is comparable to the lack of protection seen after injection with empty vector or eGFP expression vector ^20; 40^.

Importantly, even when using 2.5 times the vector concentration previously used in the Wallerian degeneration assays, expression of the NMNAT2^Q135Pfs*44^ variant was barely detectable above background by Flag immunostaining in injected neurons (Fig. 3C). In contrast, robust expression of Flag-NMNAT2^R232Q^ was observed that closely matched that of Flag-NMNAT2^WT^ (Fig. 3C). Therefore, if the truncated Flag-NMNAT2^Q135Pfs*44^ mutant retains any functionality, its inability to protect transected neurites in this assay could simply reflect very low levels of expression, whereas the failure of Flag-NMNAT2^R232Q^ to protect must instead reflect either much more rapid loss of the mutant protein after injury relative to Flag-NMNAT2^WT^, or a substantial loss of function.

### NMNAT2^Q135Pfs*44^ variant produces an unstable protein whereas NMNAT2^R232Q^ variant is slightly more stable than wild-type NMNAT2

To investigate whether the stability of either variant Flag-NMNAT2 protein is altered relative to Flag-NMNAT2^WT^, we assessed their relative rates of turnover in transfected HEK 293T cells after a protein synthesis block. Expression of the exogenous proteins was kept low to avoid saturation of the degradation machinery. Levels of Flag-NMNAT2^WT^ and Flag-NMNAT2^R232Q^ at the start of the protein synthesis block were comparable, whereas Flag-NMNAT2^Q135Pfs*44^ levels were greatly reduced (Fig. 4A, B). Flag-NMNAT2^Q135Pfs*44^ migrates at the expected size for the truncated protein but, intriguingly, Flag-NMNAT2^R232Q^ consistently migrates slightly slower than Flag-NMNAT2^WT^ (Fig. 4A). To give a more representative comparison of turnover rate of the Flag-NMNAT2^Q135Pfs*44^ mutant we increased its expression to better match starting levels of the Flag-NMNAT2^WT^ and Flag-NMNAT2^R232Q^ (Fig. 4C). In broad agreement with previous analyses, we saw almost complete loss of Flag-NMNAT2^WT^ within the 8 hour timeframe of these assays with only ∼25% remaining at 2 hours (Fig. 4A,D) ^20; 40^. In comparison, Flag-NMNAT2^Q135Pfs*44^ was undetectable on blots even at 2 hours, even from the highest starting levels (Fig 4C), whereas significantly more Flag-NMNAT2^R232Q^ was detectable at both 2 and 4 hours (Fig. 4A,D). Notably however, Flag- NMNAT2^R232Q^ was also almost completely lost by 8 hours (Fig. 4A,D).

These data indicate that Flag-NMNAT2^Q135Pfs*44^ is much less stable than Flag-NMNAT2^WT^ whereas Flag-NMNAT2^R232Q^ is modestly more stable. The reduced stability of Q135Pfs*44 Flag-NMNAT2 could thus partly explain both its lower expression level in SCG neurons and transfected HEK cells and its greatly reduced capacity to protect injured axons (above), even in the unlikely event the severely truncated protein remains functional. In contrast, the slightly increased stability of Flag- NMNAT2^R232Q^ instead suggests that its lack of axon-protective capacity is likely due to a loss of one or more other functional properties.

### NMNAT2^R232Q^ variant displays impaired NAD synthase and chaperone functions

Recombinant human NMNAT2^WT^, NMNAT2^R232Q^ and NMNAT2^Q135Pfs*44^ were obtained as His-tagged fusion proteins after bacterial expression and His-tag affinity chromatography. Typically, this yielded ∼2 mg of relatively pure recombinant protein per 0.5 L of bacterial culture for both NMNAT2^WT^ and NMNAT2^R232Q^ (Fig. 5A and 5B). Notably, the purified NMNAT2^R232Q^ was found to have NMNAT activity of just 0.51 ± 0.04 U/mg in these preparations compared to 11.2 ± 0.23 U/mg for purified NMNAT2^WT^ (Fig. 5C). In contrast, NMNAT2^Q135Pfs*44^ purifications yielded 0.1 mg or less of protein per 0.5 L of bacterial culture with a ∼22 kDa His-tagged protein, corresponding in size to NMNAT2^Q135Pfs*44^, seemingly a relatively minor component of the preparations (Fig. 5A and 5B). This is consistent with low level expression of the truncated protein in bacteria, as in mammalian cells. While the highly heterogeneous NMNAT2^Q135Pfs*44^ preparations did have detectable NMNAT activity, size-exclusion and ion exchange chromatography revealed that none of the activity was associated with the 22 kDa protein species (not shown). Instead, the invariant presence of a ∼34 kDa protein recognized by both anti-His and anti-NMNAT2 antibodies (Fig. 5A and 5B), and thus likely to be His-tagged NMNAT2^WT^, probably accounts for any activity in these preparations. Although the origin of this full-length protein remains unknown (correction of the frameshift mutation in the construct by ribosomal frameshifting or transcriptional slippage in bacteria is one possibility), this analysis suggests that the truncated NMNAT2^Q135Pfs*44^ protein is inactive as expected.

The purity of NMNAT2^R232Q^ preparations allowed us to perform further characterization that was not possible for NMNAT2^Q135Pfs*44^. NMNAT2^R232Q^ was eluted as a monomer following size-exclusion chromatography and was stable at −80 °C for months but was progressively inactivated after thawing, similar to wild type NMNAT2 ^34; 41^. There was also a linear decline of NMNAT activity for NMNAT2^R232Q^ at Mg^2+^ concentrations above 5 mM (Fig. 5D) so all subsequent assays were performed at MgCl_2_ concentrations only marginally exceeding the ATP concentration (see Methods) thus avoiding the large excess of free Mg^2+^ ions in solution usually employed for assaying NMNAT2^WT 35; 36^. Further assays revealed a relative thermal resistance and stability of NMNAT2^R232Q^ which showed a markedly higher residual activity, at least relative to its lower baseline (Fig. 5C), than NMNAT2^WT^ after 1 hour incubations at temperatures ranging from 25 °C to 47 °C (Fig. 5E) and, at ∼40 min, the activity half-life of NMNAT2^R232Q^ at 37 °C was more than twice that of NMNAT2^WT^ (Fig. 5F). Nevertheless, the optimum temperature for activity was the same for NMNAT2^WT^ and NMNAT2^R232Q^ (Fig. 5G). Crucially, however, the R232Q mutation had a profound negative effect on kinetic properties of the enzyme: *K*_cat_ was found to be reduced by ∼20-fold and the *Km* values for NMN and ATP were both increased ∼10-fold (Table 2). These striking changes predict a ∼200-fold reduced catalytic efficiency (*K*_cat_/*K*_m_) of NMNAT2^R232Q^ compared to NMNAT2^WT^ (Table 2). In fact, the loss of catalytic activity may even be greater in *vivo* where physiological concentrations of ATP (∼1 mM) and NMN (5 µM) in brain predict a 500-fold or greater reduction compared to the wild type enzyme ^42^.

**Table 2.** Kinetic parameters of human NMNAT2 WT and R232Q.

| Enzyme | Substrate | $K_m$ ($\mu\text{M}$) | $K_{cat}$ ($\text{s}^{-1}$) | $K_{cat}/K_m$ ($\text{s}^{-1}\text{M}^{-1}$) |
| --- | --- | --- | --- | --- |
| NMNAT2<sup>WT</sup> | ATP | $159.6 \pm 28.6$ | $6.84 \pm 3.21$ | $0.429 * 10^5$ |
| | NMN | $22.3 \pm 8.13$ | $6.84 \pm 3.21$ | $3.070 * 10^5$ |
| NMNAT2<sup>R232Q</sup> | ATP | $1820.9 \pm 121.1$ | $0.31 \pm 0.03$ | $0.002 * 10^5$ |
| | NMN | $178.5 \pm 10.8$ | $0.31 \pm 0.03$ | $0.017 * 10^5$ |

Together, these data suggest that both NMNAT2^R232Q^ and NMNAT2^Q135Pfs*44^ have a substantial loss of NMNAT activity. Although the R232Q mutation makes the enzyme slightly more resistant to heat denaturation *in vitro*, any increased stability is likely to be completely negated by the substantial detrimental effect it has on catalytic activity. In contrast, reduced expression/stability and impaired catalytic activity (largely predicted from the absence of key C-terminal motifs resulting from truncation) likely combine to severely impair the activity of NMNAT2^Q135Pfs*44^. The presence of only NMNAT2^R232Q^ and NMNAT2^Q135Pfs*44^ in neurons would thus be predicted to be highly limiting for NMN consumption and NAD^+^ biosynthesis in axons, thereby limiting survival.

To characterize the chaperone function, we used an in-cell luciferase refolding assay to measure the ability of NMNAT isoforms to facilitate the refolding of unfolded luciferase after heat shock ^43; 44^ (Fig. 6). Chaperones may act as “holdases” to protect their client protein from unfolding, or as “foldases” to assist the folding to the native state ^45; 46^. The in-cell luciferase-refolding assay allows the measurement of the luciferase unfolding after heat shock (red bars), as well as the luciferase refolding after recovery (green bars) ^43; 44^. We found that both NMNAT2^WT^ and NMNAT2^R232Q^ greatly protected luciferase from unfolding during heat shock, indicating strong “holdase” activity (Fig. 6, red bars). However, when “foldase” activity was analyzed, we found a remarkable loss of foldase activity specifically in NMNAT2^R232Q^ expressing cells, while NMNAT2^WT^ facilitated the refolding of luciferase after heat shock, comparable to heat shock protein 70 (Hsp70) and NMNAT3 (Fig. 6, green bars). Compared to NMNAT2^WT^ and NMNAT^R232Q^, NMNAT2^Q135Pfs*44^ did not exhibit either “holdase” or “foldase” activity, indicating a lack of stable or functional protein.

**Figure 6:**
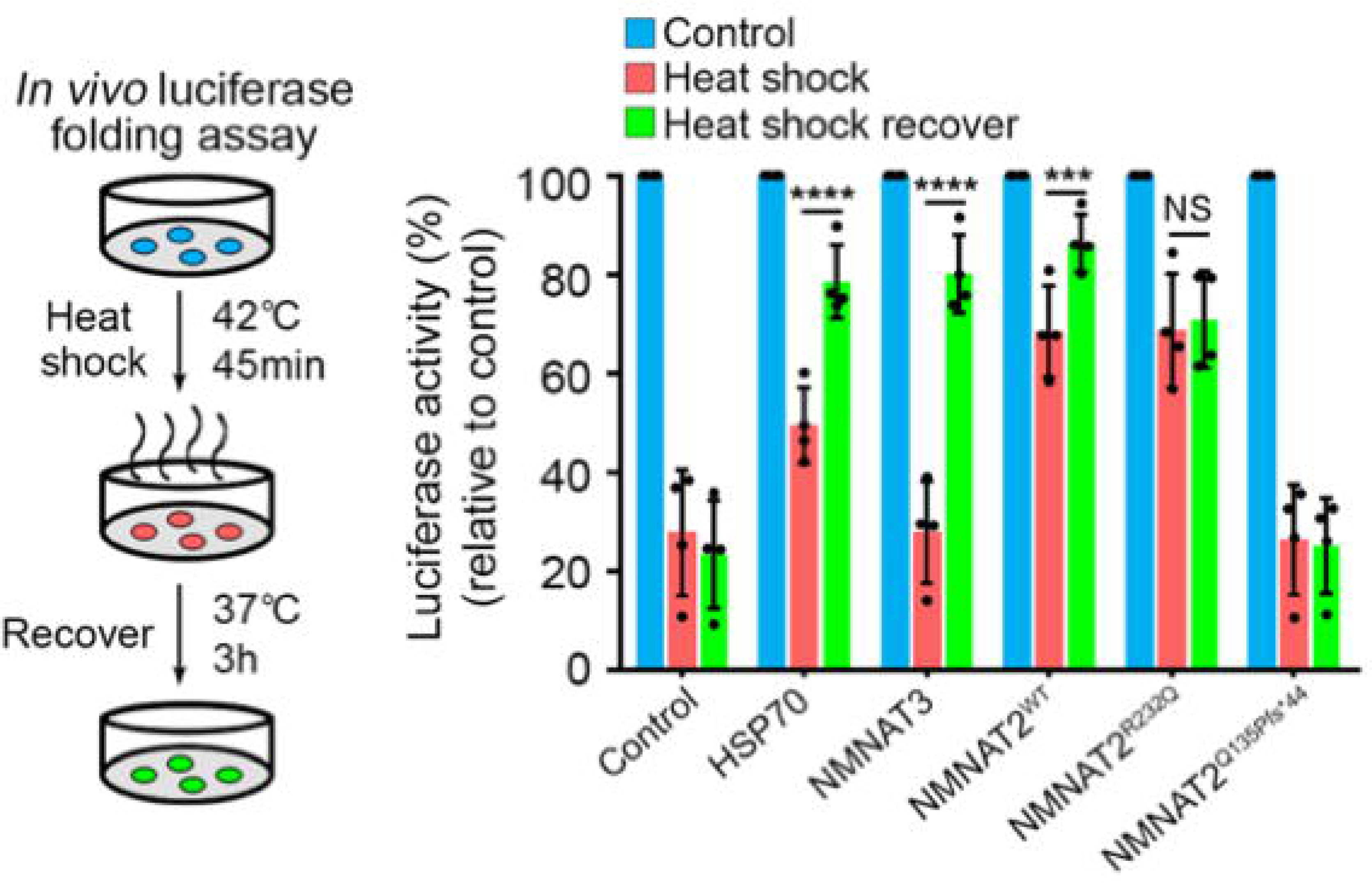
NMNAT2^R232Q^ and NMNAT2^Q135Pfs*44^ have reduced chaperone activities. HEK 293T cells were co-transfected with luciferase and one of the following plasmids: DsRed2 vector (control), Hsp70, NMNAT3, NMNAT2^WT^, NMNAT2^R232Q^, and Nmnat2^Q135Pfs*44^. At 48 hrs after transfection, protein synthesis was inhibited, and cells were subjected to heat shock at 42°C for 45 mins, and then recovered at 37 °C for 3 hours. Quantification of luciferase activity measured without heat shock (blue bars), after heat shock (red bards), and after recovery (green bars). Luciferase activity in each group was normalized to no heat shock (set to 1). All data were presented as mean ± SD, n=4. Statistical significance was established by two-way ANOVA post hoc Tukey’s multiple comparison test. ***P<0.001, ****P<0.0001, NS: not significant.

Collectively, these biochemical and cellular analyses revealed two functional consequence of the NMNAT2 mutation p.R232Q: a significant loss of enzymatic activity and a complete loss of chaperone foldase activity. Considering the essential neuronal maintenance function of NMNAT2^20; 47; 48^, disruption of both activities provides the molecular basis for compromised embryonic metabolism and neuronal development, contributing to FADS.

## DISCUSSION

The critical role of NMNAT2 in promoting axon survival in mice has been well established using *in vitro* and *in vivo* models. Declining NMNAT2 levels have been associated with a variety of neurodegenerative diseases, including Alzheimer’s disease and other tauopathies ^24; 25; 29; 47; 49^, but a direct role in human disease causation has not previously been demonstrated. Here we report two related fetuses that are both compound heterozygous for severe loss-of-function *NMNAT2* alleles. They both present with a FADS phenotype closely resembling that of homozygous null mice. Thus, for the first time, we present strong evidence that *NMNAT2* loss of function causes a human disorder.

We performed detailed molecular analyses of the *NMNAT2* variants to support our claims. We first tested the ability of the variants to protect axons in a Wallerian Degeneration model. Consistent with previous reports, the NMNAT2^WT^ construct was able to protect nearly 75% of axons from Wallerian Degeneration. However, both *NMNAT2* variants were severely compromised in their ability to delay degeneration with 10% or less of neurites remaining intact 24 hours post transection. These data argue both patient variants significantly impair NMNAT2 function in PNS axons.

We also performed *in vitro* experiments to specifically query enzymatic and chaperone functions for NMNAT2. We most conclusively demonstrated severe loss-of-function of the NMNAT2^R232Q^ variant. The strong conservation of the R232 residue across evolutionarily distant NMNAT homologues suggested that the R232 residue is functionally relevant. The invariable R residue is at the end of a β strand connecting the substrate binding domains for NMN and ATP, therefore we predicted the R232Q mutation likely affects substrates and NAD^+^ binding. Indeed, we found the R232Q substitution substantially reduces affinity for both substrates and largely abolishes NMNAT activity of the variant protein. Furthermore, R232 forms the bend between the β strand and a helix and is positioned at the surface of the protein that likely participates in the interface of protein-protein interactions. The reduced refolding activity of NMNAT2^R232Q^ is also thus consistent with reduced protein-protein interaction(s) with the cellular refolding machinery as a result of the missense mutation. Interestingly, NMNAT2^R232Q^ also consistently shows retarded migration during electrophoresis. While the R232Q missense mutation could influence migration by altering the charge and/or structural rigidity of the protein, the possibility that it might be the result of altered posttranslational modification also needs to be considered, especially in the context of the loss-of-function and increased stability of this variant.

The relative instability of the NMNAT2^Q135Pfs*^ variant protein largely precluded the same degree of functional assessment. However, this instability is probably sufficient on its own to explain the observed loss-of-function in our assays. Nevertheless, the fact that the frameshift mutation results in a truncated protein lacking its entire C-terminal half, including many residues that are critical for ATP binding, makes it extremely likely that NMNAT2^Q135Pfs*44^ will also be defective for NMNAT activity and chaperone function. Interestingly, because we expressed NMNAT2^Q135Pfs*44^ from an intronless construct in our assays, its relative instability likely reflects an increased susceptibility of the truncated peptide to direct proteolytic cleavage. However, it remains possible that nonsense mediated decay of the aberrant mRNA could also further limit expression the FADS cases.

Importantly, the FADS phenotype seen in the human patients shows broad overlap with that of *MNMAT2-deficient* mice, in particular the severely reduced skeletal muscle mass and akinesia, which are both likely due to failed peripheral innervation ^21; 22^. However, the human cases and the mouse model also show a number of notable differences. First, viability is further reduced in humans; fetal patient demise occurs at around 27 weeks in gestation whereas *MNMAT2-deficient* mice die perinatally. Second, at least one patient developed hydrocephalus in which the cortex was essentially spared. We do note *Nmnat2* shows strong expression in the CNS and PNS and therefore it is possible that *Nmnat2* deficiency is related to the hydrocephlsus ^50^. Third, the bladder is consistently distended in the mouse model but we did not identify defects in the bladder of either patient. Crucially, these differences could all be related to significantly longer gestation and longer axons in humans than mice which likely allow for more severe neurodegeneration in humans *in utero* and an increased likelihood of fetal demise ^51^. In addition, some symptoms specific to the human cases, including cystic hygroma, ascites, and edema are likely to be a consequence of the fetal demise *in utero*. Interestingly, mice nullizygous for other FADS-associated genes, such as *Dok7* and *Musk* ^52; 53^, have a phenotype remarkably similar to *MNMAT2-deficient* mice, providing additional support for a direct link between the *NMNAT2* loss-of-function alleles in these cases and their FADS presentation.

There is strong evidence in the literature from several independent groups suggesting that NMNAT2 enzymatic activity is the key activity for preventing activation of Wallerian-like axon degeneration and that enzyme dead / chaperone competent mutants broadly fail to protect axons ^13^. At the moment it is not known whether NMNAT2 chaperone function also contributes to axon protection and the finding that blocking the Wallerian degeneration pathway by removal of SARM1 “fully” rescues axon defects and survival of mice lacking NMNAT2 suggests that chaperone activity is dispensable for survival or overt health in mice, at least in the context of a relatively non-stressful home cage environment ^26; 27^. However, as we found the R232Q variant affects both chaperone and NAD synthase functions of NMNAT2, we cannot definitively exclude a critical requirement for the chaperone function in human development.

We conclude that the compound heterozygous variant *NMNAT2* alleles in the FADS cases described here encode proteins whose enzymatic and chaperone functions are both either directly or indirectly impaired and are a likely underlying cause of the disorder. As in *MNMAT2-deficient* mice, we propose that defects in PNS axon outgrowth and/or survival primarily lead to decreased innervation of the skeletal muscle in the fetuses resulting in severely reduced skeletal muscle mass. We argue *NMNAT2* may be added to the growing list of genes involved in developing or maintaining PNS innervation that have been linked to FADS or FADS-associated symptoms such as *DOK7*, *MUSK, RAPSN, ADCY6, GPR126, ECEL1, GLDN,* and *PIEZO2* ^1^. *NMNAT2* mutations should be investigated in other cases with fetal hydrops, fetal akinesia, and widespread skeletal muscle deficiency.

It will also be important to determine whether more modest *NMNAT2* loss-of-function alleles are associated with other disorders. Interestingly, in the accompanying paper [Huppke et al;], another set of patients has been identified that are homozygous for a temperature-sensitive, partial loss-of-function *NMNAT2* allele who develop a childhood-onset peripheral neuropathy phenotype. This raises the possibility of an *NMNAT2* allelic series with mutations that have a less severe effect on NMNAT2 function leading to childhood or later-onset neuropathies, rather than prenatal lethality, and/or preclinical phenotypes that predispose to adult-onset disorders.

## Supplemental Data

Two tables with more information on variants identified in whole exome sequencing.

**The authors declare no competing interests**

## Acknowledgements

Funding for the project comes from the NIH (R.W.S. R01NS085023; R.G.Z. R56NS095893), the

UK Medical Research Council grant (J.G. MR/N004582/1), the John and Lucille van Geest Foundation (M.C.) and the Taishan Scholar Project of Shandong Province, China (R.G.Z.).

