## Supplemental Figure 1 for "Severe Biallelic Loss-of-function Mutations in *Nicotinamide Mononucleotide Adenylyltransferase 2 (NMNAT2)* in Two Fetuses with Fetal Akinesia Deformation Sequence"

Supp. Table 1. Genetic Variants from whole exome sequencing of fetus II-1.

| | | Medical Review | | | | |
|---|---|---|---|---|---|---|
| | Inheritance Model Filtering | Alteration Review | Clinical Association Review | | | Candidates |
| Inheritance | Total | Total | Characterized | Clinically novel | Total | Total |
| Autosomal Dominant | 3(4) | 0(0) | 0(0) | 0(0) | 0(0) | 0(0) |
| Autosomal Recessive | 7(17) | 1(2) | 0(0) | 1(2) | 1(2) | 1(2) |
| X-linked Recessive | 0(0) | 0(0) | 0(0) | 0(0) | 0(0) | 0(0) |
| X-linked Dominant | 0(0) | 0(0) | 0(0) | 0(0) | 0(0) | 0(0) |
| Autosomal Dominant (reduced penetrance) | 18(22) | 8(8) | 0(0) | 0(0) | 0(0) | 0(0) |
| X-linked (reduced penetrance) | 0(0) | 0(0) | 0(0) | 0(0) | 0(0) | 0(0) |
| All Models | 28 (43) | 9(10) | 0(0) | 1(2) | 1(2) | 1(2) |

Supp. Table 2. Alternative variants identified in fetus II-1 whole exome sequencing.

| Gene | Locus | RefSeq ID | Alteration | OMIM | Proband | Father | Mother | Genome1k | PolyPhen | Sift |
|---|---|---|---|---|---|---|---|---|---|---|
| ***CACNA1S*** | 1:201020165 | NM_000069 | c.4060A>T p.T1354S | Thyrotoxic periodic paralysis, susceptibility to, 1<br>Malignant hyperthermia susceptibility 5<br>Hypokalemic periodic paralysis, type 1 | +/- | +/- | -/- | 0.86% | benign | tolerated |
| ***COL6A1*** | 21:47404305 | NM_001848 | c.350T>C p.V117A | Ossification of the posterior longitudinal spinal ligaments | +/- | +/- | -/- | N/A | probably damaging | unknown |
| ***COL7A1*** | 3:48602833-48602846 | NM_000094 | c.8524_8527+10DELGAAGGTGAGGACAG | Transient bullous of the newborn<br>Toenail dystrophy, isolated<br>Epidermolysis bullosa, pretibial<br>Epidermolysis bullosa dystrophica, AR<br>Epidermolysis bullosa dystrophica, AD | +/- | +/- | -/- | N/A | N/A | N/A |
| ***DSP*** | 6:7580795 | NM_004415 | c.4372C>G p.R1458G | Skin fragility-woolly hair syndrome<br>Keratosis palmoplantaris striata II<br>Dilated cardiomyopathy with woolly hair and keratoderma<br>Arrhythmogenic right ventricular dysplasia 8 | +/- | +/- | -/- | 0.91% | benign | tolerated |
| ***EFCAB5*** | 17:28407871-28407874 | NM_198529 | c.3298_3301DUPAATG | N/A | +/- | -/- | +/- | N/A | N/A | N/A |
| ***HCN4*** | 15:73616159 | NM_005477 | c.2275G>A p.V759I | Brugada syndrome 8<br>Sick sinus syndrome 2 | +/- | +/- | -/- | 1.76% | benign | tolerated |
| ***MYO15A*** | 17:18051447 | NM_016239 | c.6614C>T p.T2205I | Deafness, autosomal recessive 3 | +/- | -/- | +/- | 1.61% | probably damaging | deleterious |
| ***PKHD1*** | 6:51908433 | NM_138694 | c.2811G>A p.W937* | Polycystic kidney and hepatic disease | +/- | -/- | +/- | N/A | N/A | N/A |
| ***PKLR*** | 1:155261570 | NM_000298 | c.1595G>A p.R532Q | Adenosine triphosphate, elevated, of erythrocytes | +/- | +/- | -/- | N/A | probably damaging | deleterious |
| ***RB1*** | 13:49033812 | NM_000321 | c.1961-12T>C | Small cell cancer of the lung, somatic<br>Retinoblastoma, trilateral<br>Osteosarcoma, somatic<br>Bladder cancer, somatic | +/- | +/- | -/- | 0.59% | N/A | N/A |
| ***SERPINA1*** | 14:94844947 | NM_000295 | c.1096G>A p.E366K | Pulmonary disease, chronic obstructive, susceptibility to<br>Hemorrhagic diathesis due to 'antithrombin' Pittsburgh<br>Emphysema-cirrhosis, due to AAT deficiency<br>Emphysema due to AAT deficiency | +/- | -/- | +/- | 4.12% | probably damaging | deleterious |
